# Progressive enhancement of kinetic proofreading in T cell antigen discrimination from receptor activation to DAG generation

**DOI:** 10.1101/2021.11.10.468056

**Authors:** Derek Britain, Orion Weiner

## Abstract

T cells use kinetic proofreading to discriminate antigens by converting small changes in antigen binding lifetime into large differences in cell activation, but where in the signaling cascade this computation is performed is unknown. Previously, we developed a light-gated immune receptor to probe the role of ligand kinetics in T cell antigen signaling. We found significant kinetic proofreading at the level of the signaling lipid diacylglycerol (DAG) but lacked the ability to determine where the multiple signaling steps required for kinetic discrimination originate in the upstream signaling cascade (Tischer and Weiner, 2019). Here we uncover where kinetic proofreading is executed by adapting our optogenetic system for robust activation of early signaling events. We find the strength of kinetic proofreading progressively increases from Zap70 recruitment to LAT clustering to downstream DAG generation. These data suggest a distributed kinetic proofreading mechanism, with proofreading steps both at the receptor and at downstream signaling events. Leveraging the ability of our system to rapidly disengage ligand binding, we measure slower reset rates for downstream signaling events. Our observations of distributed kinetic proofreading and slowed resetting of downstream steps suggest a basis of cooperativity between multiple active receptors with implications in tissue homeostasis, autoimmunity, and immunotherapy off-target effects.

## Introduction

Proper antigen discrimination is a cornerstone of the adaptive immune system. Failure of T cells to activate in response to foreign antigen can enable pathogens to invade the body undetected (Charles A Janeway et al., 2001; Horst et al., 2011). Conversely, improper T cell activation against self-antigen causes autoimmune disorders (Dornmair et al., 2003). When a T cell contacts an antigen-presenting cell (APC), it can detect the presence of 1-10 molecules of foreign antigen (Christinck et al., 1991; Demotz et al., 1990; Kimachi et al., 1997; Sykulev et al., 1996), despite the much greater abundance (100,000x or more) of self antigen (Bhardwaj et al., 1993; Cohen et al., 2003; Irvine et al., 2002; Unternaehrer et al., 2007). These observations indicate that T cells must use parameters other than the number of ligand-bound receptors to distinguish self from foreign antigen (Daniels et al., 2006; M. M. Davis et al., 1998; Gascoigne et al., 2001; Germain & Stefanová, 1999).

One attractive model for how T cells discriminate antigen is the kinetic proofreading model (McKeithan, 1995; Ninio, 1975), where only antigens that continuously bind a TCR for a sufficient duration activate the T cell (**Fig. 1A**). Kinetic proofreading postulates that antigen binding begins a series of slow successive biochemical events that must progress to completion before activating the T cell. Only antigen that stays bound to the TCR long enough for the completion of all events is stimulatory to the T cell (**Fig. 1B**). If the antigen unbinds the TCR before all events are complete, the system resets back to the ground state (**Fig. 1C)**. While it is commonly accepted that T cells use kinetic proofreading for antigen discrimination (Coombs & Goldstein, 2005; McKeithan, 1995), we do not know which steps in the antigen signal transduction cascade enable the strong kinetic proofreading observed in T cell activation.

**Figure 1.**
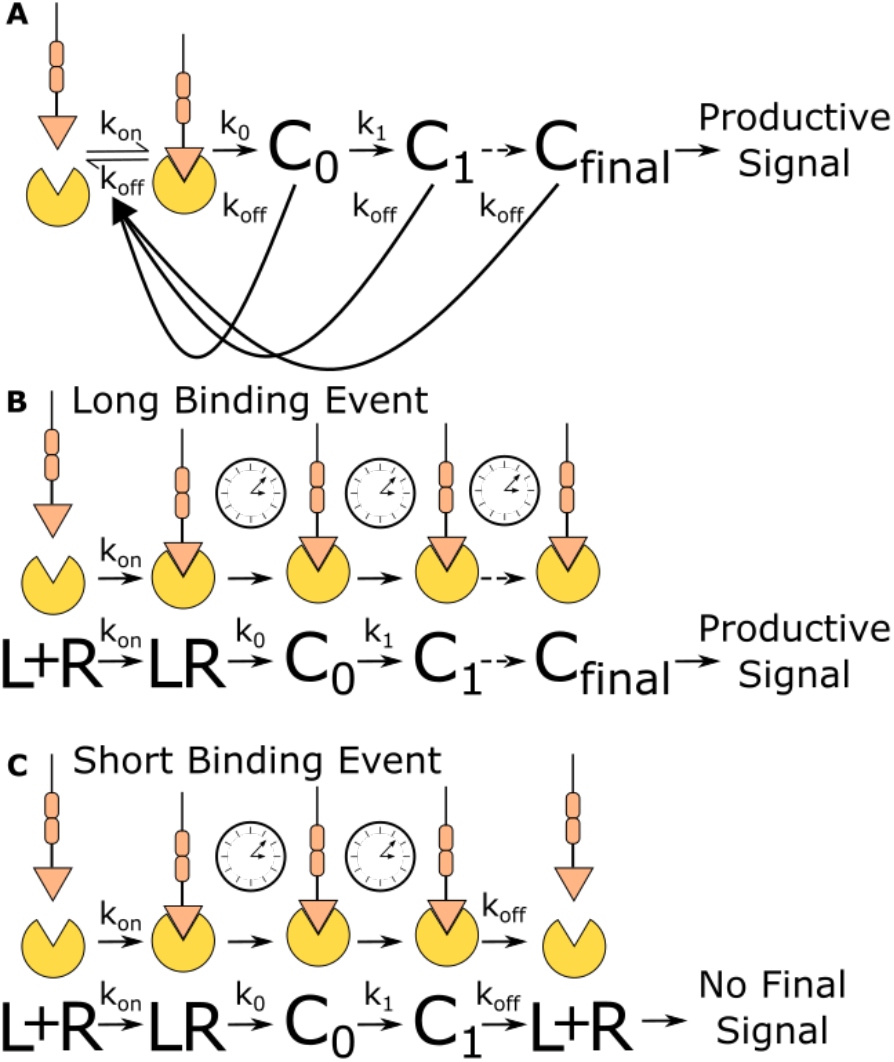
Kinetic proofreading discriminates binding events by half-life. (**A**) The kinetic proofreading model consists of a chain of slow sequential events (C_i_ with rates k_i_) that begin once the ligand binds the receptor (k_on_). The chain of events must be completed before a productive signal is communicated to the cell. When the ligand unbinds the receptor (k_off_), all accumulated signaling intermediate (C_i_) reset back to the ground state. Only binding events that persist long enough to form the active terminal signaling complex (C_final_) result in cell activation. **(B)** Long-lived binding events remain bound to the receptor for a sufficient duration to complete all kinetic proofreading steps, resulting in a productive signal. (**C**) Short-lived binding events unbind the receptor before the kinetic proofreading chain is completed and fail to fully stimulate the antigen signaling cascade.

Kinetic proofreading can convert small differences in ligand binding lifetime into large differences in cellular output. The number of proofreading steps sets the strength of a proofreading system, with stronger proofreading systems requiring a higher number of proofreading steps. Ten steps of kinetic proofreading would roughly amplify a 2 fold difference in ligand binding lifetime into a 1,000 fold difference in output. Kinetic proofreading allows T cells to ignore numerous short-lived self antigen binding events, while activating in response to rare long-lived foreign antigen binding events (M. M. Davis et al., 1998; Gascoigne et al., 2001; Germain & Stefanová, 1999).

Previously, our group developed a light-gated immune receptor that we used to probe the role of ligand kinetics in T cell antigen signal transduction (**Fig. 2A**). We found evidence of proofreading at the level of DAG generation, while detecting no proofreading at the recruitment of Zap70 to the activated receptor (Tischer & Weiner, 2019). The strength of proofreading measured at DAG generation indicated multiple upstream proofreading events. Our optogenetic system requires a light-insensitive mode of cell adhesion to ensure reliable measurement of signaling activity readouts. However, our previous adhesion method was not compatible with robust cellular activity readouts upstream of DAG generation, preventing us from identifying where the multiple upstream proofreading events originate.

**Figure 2.**
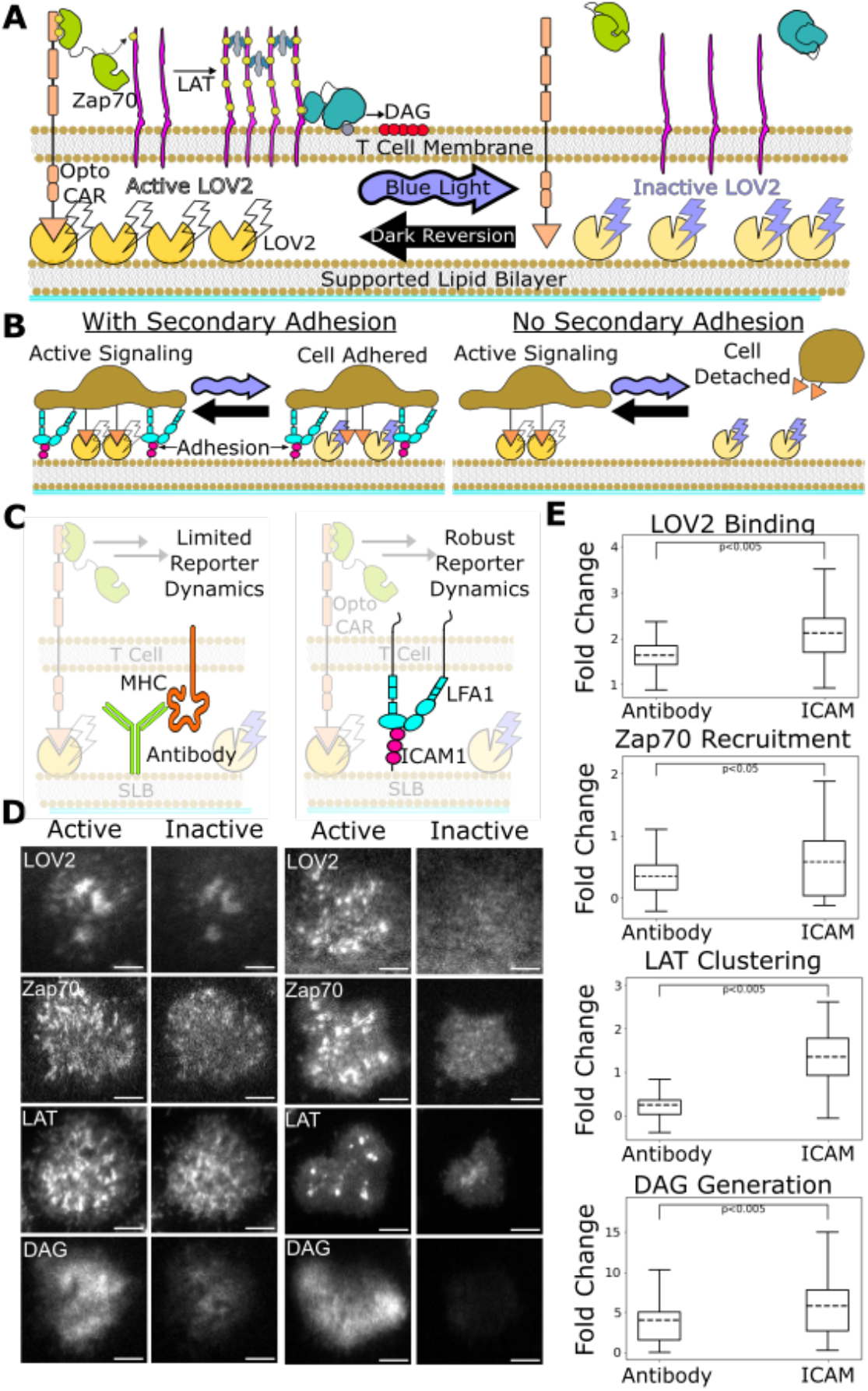
ICAM-1 adhesion improves robustness of proximal signaling live-cell biosensors. **(A)** With our optogenetic system, we can control the concentration and lifetime of ligand-receptor interactions with blue light. A supported lipid bilayer (SLB) functionalized with the light-sensitive protein LOV2 creates an artificial antigen-presenting surface. Jurkat cells expressing zdk-CAR bind dark-state LOV2 and activate antigen signaling. Blue light interrupts LOV2-CAR binding events and terminates signaling. **(B)** Our assay requires a secondary light-insensitive mode of cell adhesion to maintain segmentable and trackable cell footprints throughout an experiment. Cells adhere to the SLB through LOV2 while actively signaling. However, when signaling is inactivated by blue light, cells detach from the bilayer if no secondary method of constant adhesion exists. **(C)** In our previous work, we adhered cells to the SLB through biotinylated beta2 microglobulin antibodies against the cells’ MHC class1 (left). This adhesion technique limited the spatiotemporal dynamics of many biosensors, including the recruitment of ZAP70 and the clustering of LAT. We now adhere cells to the bilayer through native LFA-1 to ICAM-1 integrin signaling (right). **(D)** Representative TIRF images of LOV2 binding, Zap70 recruitment, LAT clustering, and DAG generation during ligand engagement and after two minutes of strong inactivation of the Zdk-CAR under antibody adhesion (left) versus ICAM-1 adhesion (right) (scale bar = 5um). (**E**) SLB adhesion through ICAM-1 improved the spatiotemporal dynamics of each biosensor. Boxplot of the maximum fold-change in LOV2 binding, Zap70 recruitment, LAT clustering, and DAG generation between active and inactive Zdk-CAR signaling states under antibody adhesion and ICAM-1 adhesion. Dashed line indicates population mean, each plot represents reporter change of approximately n=120 cells, independent t-test p values shown.

Here we improve our assay by incorporating the native integrin ICAM-1 as our light-insensitive mode of adhesion, which improves the robustness of our previous biosensors and enables the use of new biosensors (Bromley & Dustin, 2002). With integrin adhesion, we now detect kinetic proofreading in receptor activation upstream of the recruitment of Zap70. We measure stronger upstream kinetic proofreading at the clustering of the scaffold protein LAT, suggesting proofreading steps between Zap70 recruitment and LAT cluster formation. Furthermore, we measure even stronger kinetic proofreading at DAG generation, suggesting additional proofreading steps after the initial formation of LAT clusters. We also find that LAT clusters reset slower than Zap70 recruitment upon unbinding antigen. Our results suggest a kinetic proofreading system that starts at the level of receptor activation and continues across multiple spatially segregated signaling complexes, with downstream signaling complexes resetting slower than receptor-level signaling complexes. Such a system would allow high concentration of antigens with intermediate binding lifetimes to activate T cells, while still responding to rare long-binding antigens and filtering out short-binding antigens, which is consistent with TCR cooperativity postulated in tissue homeostasis and autoimmunity (Cameron et al., 2013; Goyette et al., 2020; Korem Kohanim et al., 2020; Lin et al., 2019; Pettmann et al., 2021; Wang et al., 2020).

## Results

### Expanding suite of optogenetically-controlled T cell biosensors by optimizing cell adhesion system

Previously we developed a light-gated immune receptor (zdk-CAR) to probe T cell antigen signaling output as a function of both receptor occupancy and average ligand-binding half-life (**Fig 2A**) (Tischer & Weiner, 2019). In the dark, cells expressing zdk-CAR bind to purified LOV2 ligands functionalized on a supported lipid bilayer (SLB) and activate antigen signal transduction. Blue light interrupts LOV2 binding events and antigen signaling. If we provide LOV2 as the only means of cell interaction with the bilayer, cells unbind from the bilayer upon exposure to blue light (**Fig. 2B**). This complicates our analysis, which requires exposing cells to multiple doses of blue light to build up a dose-response curve. Therefore we must also include a light-insensitive mode of adhering the cells to the bilayer.

In our previous work, we adhered cells to the SLB using an antibody against their own MHC (Abcam ab21899) (**Fig. 2C, left**). However, this method was not compatible with sensitive detection of some cellular activity readouts, particularly the most proximal ones like Zap70 recruitment. In this work we improve our assay by adhering the cells to the SLB through native ICAM-1 integrin adhesion, which generates large cellular footprints while being compatible with a larger suite of antigen signaling biosensors.

We functionalized our SLB with human ICAM-1 to better mimic the T cell / APC interaction (**Fig. 2C, right**) (Dustin et al., 2007). We first validated Zap70 recruitment, LAT clustering, and DAG generation biosensors expressed in Jurkat cells on ICAM-1 functionalized bilayers (**Fig. 2D**). Jurkat cells stably expressing zdk-CAR were exposed to SLBs functionalized with purified human ICAM-1-His and alexa fluor 488 labeled LOV2. To increase integrin binding, we changed our imaging buffer to a modified HBSS (mHBSS) buffer with higher Mg^++^ concentrations (Labadia et al., 1998). Compared to our previous antibody-based adhesion, Jurkats on ICAM-1 functionalized bilayers released a greater percentage of the bound LOV2 ligand upon blue-light illumination. Adhesion through ICAM-1 also increased the dynamic range of the Zap70, LAT, and DAG biosensors (**Fig. 2E, Videos 1-4**). ICAM-1 adhered cells also displayed receptor/Zap70 and LAT clustering spatial patterns similar to those observed by others using native receptor ligands on SLBs (Balagopalan et al., 2015; Chakraborty & Weiss, 2014; Kumari et al., 2015).

### Quantifying kinetic proofreading in Zap70 recruitment, LAT clustering, and DAG generation under ICAM-1 adhesion

With our improved ICAM-1 adhesion method and expanded suite of biosensors, we applied our previous assay for measuring the strength of upstream kinetic proofreading to the signaling events of Zap70 recruitment, LAT clustering, and DAG generation. In our assay, we independently probe the effect receptor occupancy or ligand binding half-life on a given T cell intracellular signal (Tischer & Weiner, 2019). By titrating the intensity of LOV2 stimulating light, we modulate the lifetime of receptor-LOV2 binding events and the downstream antigen signaling response. By repeating the experiment at different levels of LOV2 ligand on the bilayer, we can uncouple the effects of ligand occupancy from ligand-binding half-life on cell activation.

After exposing zdk-CAR expressing cells to a LOV2 and ICAM-1 functionalized SLB, we illuminate the cells with an intensity of blue light with a known ligand binding half-life (**Fig. 3A**). After three minutes of illumination (when a steady state is reached), we image the amount of bound fluorescently tagged LOV2 accumulated underneath each cell using a long exposure image (O’Donoghue et al., 2013). We also image the output intensity of the signaling biosensor. We then reset the system with a two minute pulse of intense blue light to terminate all signaling before proceeding to expose the cells to the next intensity of blue light. We repeat the experiment on bilayers functionalized with different densities of LOV2 to sample a wider range of occupancies. Following this experimental protocol, we build up a dataset of signaling output as a function of both ligand occupancy and ligand half-life (**Fig. 3B**).

**Figure 3.**
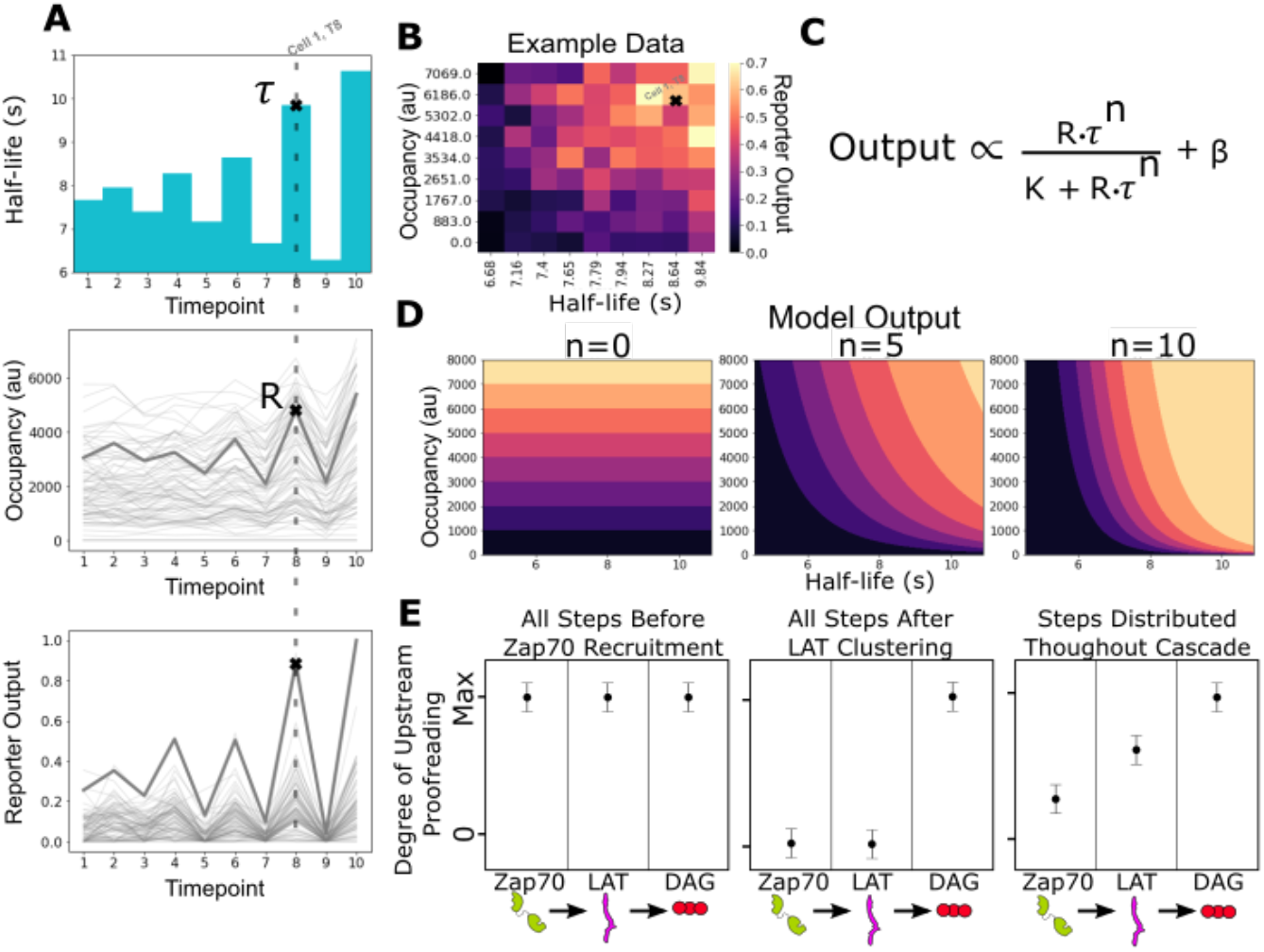
Measurement of receptor occupancy, binding half-life, and signaling output for evaluating proofreading strength. **(A)** After adhering the cells to the functionalized SLBs, we expose them to a series of three minutes blue-light conditions with known average ligand binding half-lives. At the end of each half-life condition, we measure every cell’s steady-state reporter output and receptor occupancy (example cell highlighted at time point eight). Following this protocol, we measure every cell’s reporter output (bottom) as a function of receptor occupancy (middle) and average ligand binding half-life (top). **(B)** After normalizing each cell to its average basal reporter activity, the data from multiple time courses with different ligand densities are aggregated, and the reporter output values are normalized to the 90th percentile cell in the dataset. **(C)** The dataset is then fit to a model of the expected output of a kinetic proofreading signaling system. In the model, the expected signaling output is proportional to the ligand occupancy (**R**) and ligand binding half-life (**τ**) raised to the number of strong proofreading steps (**n**) upstream of that signaling step. The magnitude of n provides a relative measure of the strength of kinetic proofreading between ligand binding and a given signaling step. K is the amount of upstream signal for half-maximal response, and β is basal signaling in the pathway (**D**) Expected output of models with zero, five, and ten kinetic proofreading steps. As the value of **n** increases, the dependence of output on ligand half-life increases, while the dependence on receptor occupancy decreases. A system with no proofreading responds only to receptor occupancy, while a system with a high degree of proofreading (n=10) is insensitive to numerous short binding events while fully responding to very few long binding events. (**E**) Anticipated results for different scenarios of proofreading step distribution.

Next we fit our data to a simple model of the expected output from a kinetic proofreading signaling system **(Fig. 3C)**. In the model, signaling output is a function of receptor occupancy (**R**) multiplied by ligand binding half-life (**τ**) raised to the number of strong proofreading steps (**n**). The fit number of proofreading steps indicates how much a reporter’s output depends on binding half-life compared to receptor occupancy. In a system with no proofreading steps, the output only depends on receptor occupancy, and binding half-life becomes irrelevant (**Fig. 3D, left**). A system’s output with a moderate number proofreading steps (e.g. five steps) depends on both receptor occupancy and the half-life of ligand binding (**Fig. 3D, middle**). A large number of proofreading steps (e.g. ten steps) results in a system whose output is dominated by ligand binding half-life and is relatively insensitive to receptor occupancy (**Fig. 3D, right**).

While certain assumptions of the model, such as equivalent strength of all proofreading steps, make the fit value of **n** unlikely to represent the absolute number of upstream biochemical proofreading steps, we can compare the measured values of **n** between signaling events to understand how the strength of proofreading changes as we progress through the cascade. A higher measured value of **n** at an event suggests a stronger degree of upstream kinetic proofreading. If we subtract the fit **n** value of a downstream event from an upstream event, we get the measured increase in proofreading strength between the two events. For example, if we have the network C_0_ → → C_x_ and measure **n**=3 at C_0_ and **n**=5 at C_x_, subtracting C_0_ from C_x_ results in a proofreading strength increase of 2, indicating additional kinetic proofreading between C_0_ and C_x._

Comparing the measured degree of proofreading at multiple known signaling events allows us to bracket where kinetic proofreading steps exist **(Fig. 3E**). If all proofreading steps occured between ligand binding and Zap70 recruitment, we would measure the same degree of upstream proofreading at Zap70 recruitment, LAT clustering, and DAG generation (**Fig. 3E, left**), as there would be no additional increase in proofreading strength beyond Zap70. If all proofreading steps occured after LAT clustering and before DAG generation, we would measure no degree of upstream proofreading at Zap70 recruitment or at LAT clustering, and see a large jump in the degree of upstream proofreading at DAG generation (**Fig. 3E, middle**). If proofreading steps were distributed from before Zap70 recruitment to after LAT clustering, we would measure a non-zero degree of upstream proofreading at Zap70 recruitment with an increasing degree of upstream proofreading as we progress down the cascade to LAT clustering and DAG generation (**Fig 3E, left**). As we will show, this last model is most consistent with our data.

### Kinetic proofreading steps exist between ligand-receptor binding and Zap70 recruitment

In our previous work, we failed to detect kinetic proofreading in Zap70 recruitment (Tischer & Weiner, 2019). Using ICAM-1 functionalized bilayers, we now measure moderate amounts of kinetic proofreading in the recruitment of Zap70 to activated receptors. Zap70 recruitment correlated with both receptor occupancy (**ρ** = 0.42) and ligand binding half-life (**ρ** = 0.49) (**Fig. 4S1A**). Our model fit a moderate degree of proofreading at Zap70 recruitment across three datasets (**n** = 4.5 +/- 0.4) (**Fig. 4A**). These data suggest the existence of kinetic proofreading steps between ligand binding and Zap70 recruitment.

**Figure 4.**
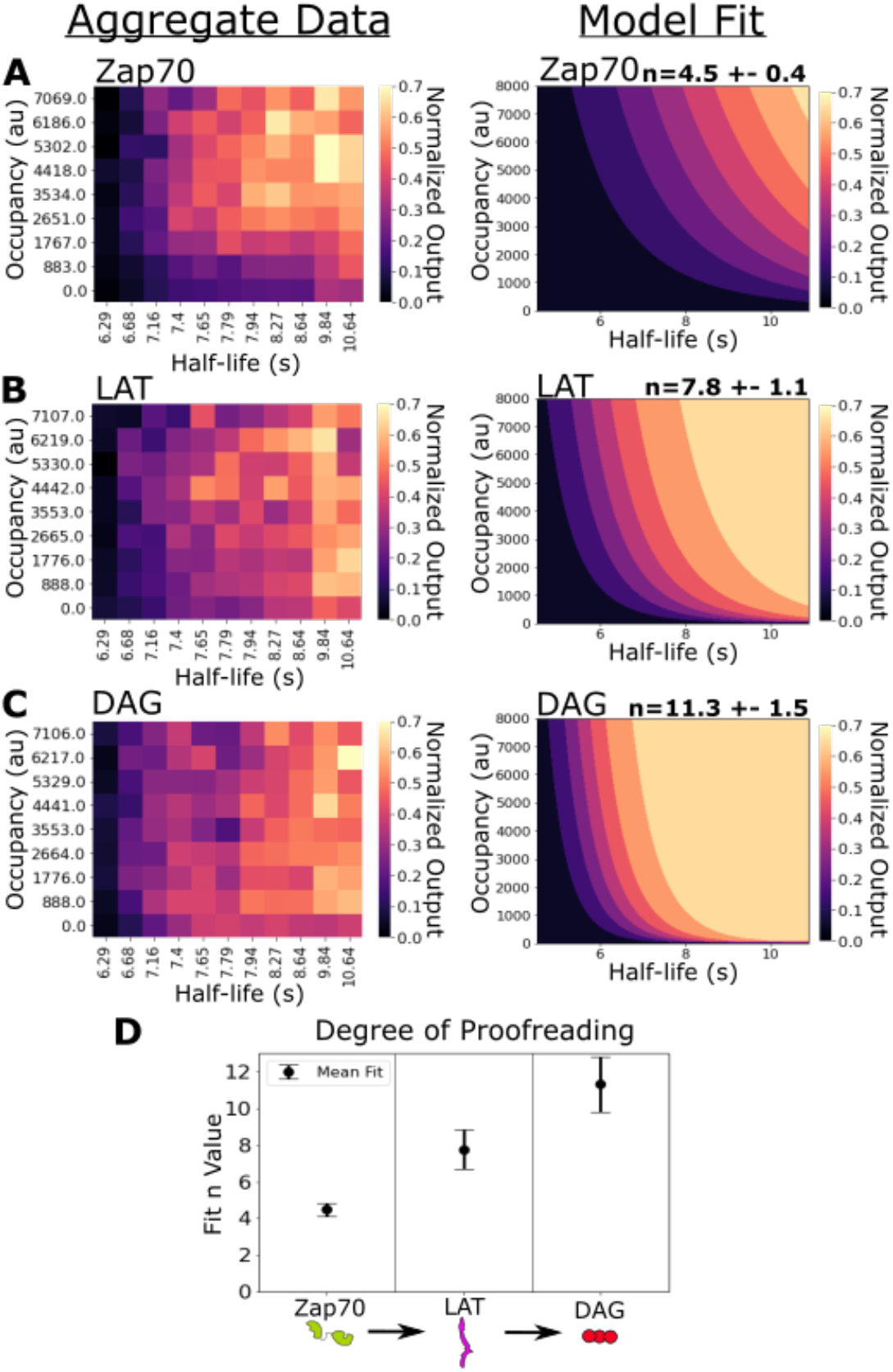
Kinetic proofreading starts upstream of Zap70 recruitment and progressively increases in strength at LAT clustering and DAG generation. (**A**) Collected data of Zap70 recruitment as a function of receptor occupancy (y-axis) and ligand half-life (x-axis) (left). Zap70 recruitment best fits a model of kinetic proofreading with a strength of upstream proofreading of **n** = 4.5 +- 0.4 (right). (**B**) Collected data of LAT clustering as a function of receptor occupancy (y-axis) and ligand half-life (x-axis) (left). LAT clustering best fits a model of kinetic proofreading with a strength of upstream proofreading of **n** = 7.8 +- 1.1 (right). (**C**) Collected data of DAG generation as a function of receptor occupancy (y-axis) and ligand half-life (x-axis) (left). DAG generation best fits a model of kinetic proofreading with a strength of upstream proofreading of **n** = 11 +- 1.5 (right). (**D**) The mean model fit n-values (+/- 1 std) for each biosensor across three biological replicates. The fit n-values continue to increase from Zap70 recruitment to LAT cluster to DAG generation. These results suggest the existence of kinetic proofreading steps between ligand-binding and Zap70 recruitment, between the recruitment of Zap70 and the formation of a LAT cluster, and between the initial formation of a LAT cluster and the generation of DAG from that cluster.

Previously, we measured Zap70 with no light-insensitive adhesion to the bilayer, as antibody adhesion inhibited Zap70 reporter dynamics (Tischer & Weiner, 2019). With no additional adhesion, only actively signaling cells adhere to the bilayer. In the absence of adhesion beyond LOV2, we potentially missed sampling high occupancy, short half-life regimes with little Zap70 recruitment, as those cells failed to adhere to the bilayer. If we filter out the short half-life conditions of our new Zap70 data (with ICAM-1 adhesion) and refit our kinetic proofreading model, we measure a similarly low proofreading result as we previously observed in the absence of adhesion beyond LOV2 (**n**=0.7 +/- 0.3) (**Fig. 4S2**). With ICAM-1 adhesion we likely capture a more complete dataset of Zap70 recruitment, improving our measurement of upstream proofreading.

### Evidence for kinetic proofreading steps between Zap70 recruitment and LAT clustering, with further steps between LAT clustering and DAG generation

Next we measured the strength of kinetic proofreading at the levels of LAT clustering and DAG generation. LAT clustering showed an increased dependency on binding half-life (**ρ**=0.54) and a decreased dependence on receptor occupancy (**ρ**=0.21) compared to Zap-70 recruitment (**Fig. 4S1B**). Our proofreading model fits our LAT clustering data with a higher degree of proofreading (**n**=7.8+/-1.1) compared to Zap70 recruitment (**Fig. 4B**). The stronger degree of kinetic proofreading at LAT clustering versus Zap70 recruitment suggests additional steps of kinetic proofreading between the recruitment of Zap70 to phosphorylated ITAMs and the formation of LAT clusters. The generation of DAG also depended heavily on ligand half-life (**ρ**=0.52), while depending the least on occupancy (**ρ**=0.18) (**Fig. 4S1C)**. Our model fits the highest degree of kinetic proofreading at DAG generation (**n**=11.3 +/-1.5), suggesting further kinetic proofreading steps beyond initial LAT clustering and upstream of DAG generation (**Fig. 4C**). The sequential increase in the degree of kinetic proofreading progressing down the antigen signaling cascade suggests steps contributing to kinetic proofreading exist throughout the cascade (**Fig. 4D**).

### LAT clusters reset more slowly than Zap70 clusters upon ligand disengagement

Kinetic proofreading requires all signaling intermediates to reset upon ligand unbinding. While it is often assumed signaling intermediates reset at similar rates, slower reset rates for downstream signaling intermediates could improve antigen discrimination (McKeithan, 1995). Our zdk-CAR deactivates with blue light, giving us the unique ability to unbind all antigen binding events synchronously. After measuring additional kinetic proofreading steps downstream of the activated receptor, we used our Zdk-CAR system to measure the off-rate of bound LOV2, recruited Zap-70, and clustered LAT following acute antigen unbinding (**Fig 5A)**.

**Figure 5.**
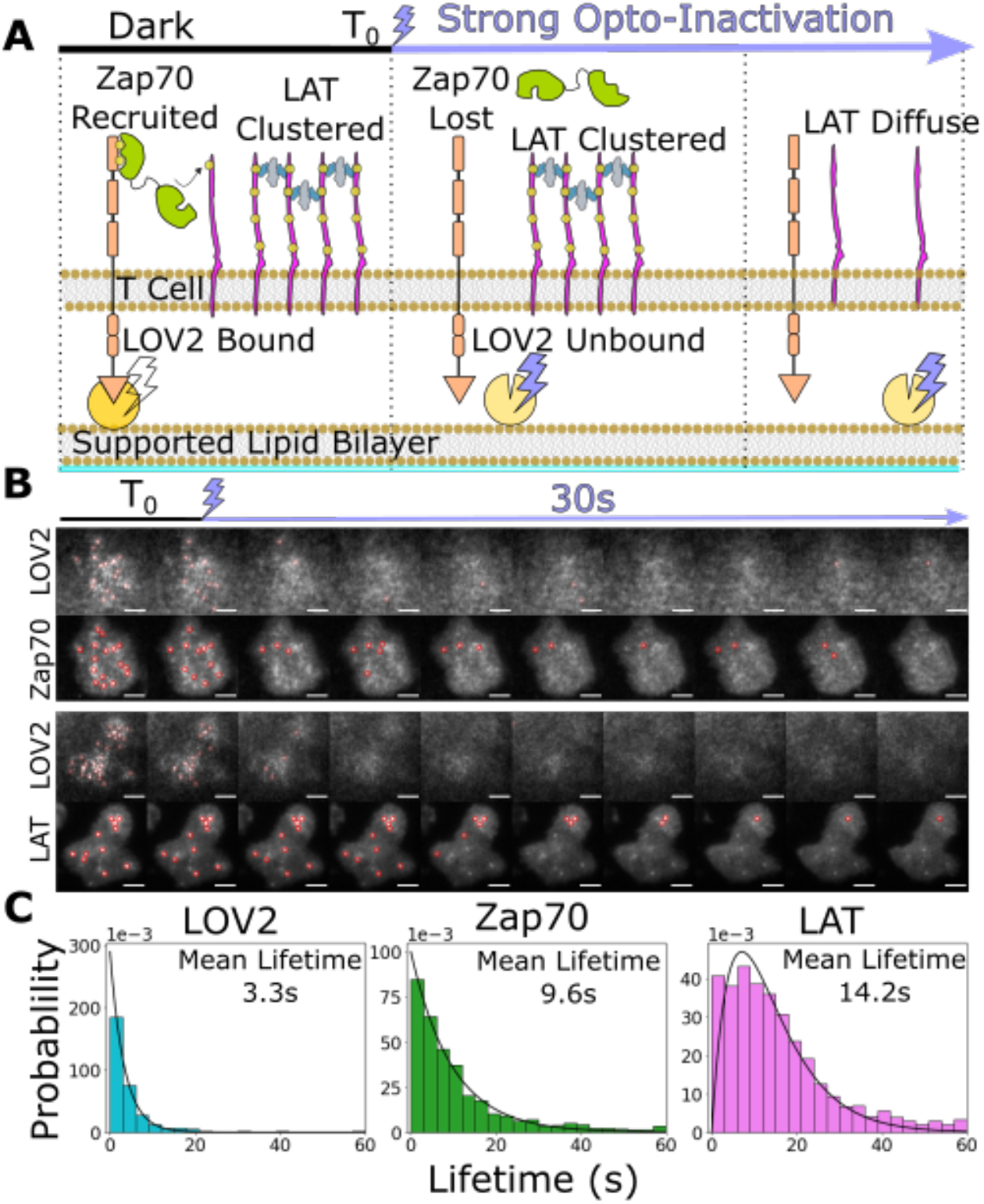
Downstream signaling complexes reset slower than active receptors upon ligand unbinding. **(A)** Schematic of reset experiment. After synchronously unbinding all receptors with intense blue light, we measure the reset rate of LOV2 unbinding, Zap70 loss, and LAT cluster dissociation. **(B)** Representative cells of Zap70 and LAT upon receptor inactivation with respective LOV2 images. Cells expressing either Zap70-Halo (top) or LAT-Halo (bottom) were allowed to activate for 3 minutes before acute inactivation with intense blue light for one minute and subsequently imaged every 3 seconds. The lifetimes of subcellular clusters of LOV2, Zap70, and LAT were tracked with the ImageJ TrackMate plugin. **(C)** The lifetime distributions of tracked clusters after receptor inactivation. The lifetime distributions of LOV2 unbinding and Zap70 dissociation fit exponential distributions with mean lifetimes of 3.3 and 9.3 seconds respectively, suggesting a one-step reset process. The LAT declustering lifetime distribution fit an Erlang distribution of a 2 step process and a mean lifetime of 14.2 seconds. Fitting of cluster intensity over time with exponentials gave comparable results for all three reporters (**Fig. 5S1**).

After allowing cells to activate on LOV-AF488 and ICAM-1-HIS functionalized bilayers for three minutes, we acutely unbound all LOV2 ligands with intense blue light while imaging LOV2 and Zap70 or LAT biosensors. We manually segmented cells in ImageJ and tracked individual sub-cellular clusters of each biosensor using the TrackMate plugin (Tinevez et al., 2017) to sample their lifetime distributions (**Fig. 5B, Videos 5 and 6**). We assumed the loss of each biosensor consisted of one or more Poisson steps. A single step mechanism results in exponentially distributed lifetimes, while a multistep process creates an Erlang lifetime distribution, where the integer shape parameter estimates the number of steps (Huang et al., 2019). We found the lifetimes of bound LOV2 and recruited Zap70 best fit exponential distributions with mean lifetimes of 3.3 s and 9.3 s, respectively. However, the lifetimes of LAT clusters best fit a two step Erlang distribution (shape= 2) with a mean lifetime of 14.2 s (**Fig. 5C**). Upon ligand unbinding, recruited Zap70 and clustered LAT likely reset through different processes, with LAT resetting through a slower multi-step process. These data open the possibility that a LAT cluster could survive a momentary ligand unbinding event (loss of Zap70) and survive long enough to integrate multiple ligand binding events, as observed in primary T cells on SLBs with native ligand (Lin et al., 2019).

## Discussion

Kinetic proofreading enables T cells to discriminate self-from cognate-antigens by converting small changes in antigen binding half-life to large changes in cell activation. Amplifying small changes in binding half-life requires a chain of multiple slow biochemical events that separate short binding events from long binding events. Where these events exist in the antigen signaling cascade is not fully understood.

In this study, we improved our optogenetic proofreading assay with ICAM-1 adhesion and measured the dependency of Zap70 recruitment, LAT clustering, and DAG generation on antigen binding half-life and receptor occupancy. We now find evidence for kinetic proofreading in antigen signal transduction as early as Zap70 recruitment to phosphorylated receptor ITAMs. We measured greater kinetic proofreading at the clustering of the scaffold protein LAT, and the highest degree of kinetic proofreading at the generation of the signaling lipid DAG (**Fig. 4D**). We also measure the reset rate of Zap70 and LAT upon ligand-unbinding. Our findings suggest that the kinetic proofreading underlying T cell antigen discrimination spans multiple membrane-associated signaling complexes (Yi et al., 2019).

After a receptor binds an antigen, many events must occur before Zap70 is recruited to the receptor’s ITAMs. Many strong candidates exist for the mediators of this kinetic processing (Chakraborty & Weiss, 2014). Ligand binding excludes the bulky phosphatase CD45 that would otherwise rapidly dephosphorylate the ITAMs (S. J. Davis & van der Merwe, 2006; Springer, 1990). The receptor itself could undergo mechanical conformational changes making it more accessible for phosphorylation (Kim et al., 2009, 2010). An active molecule of the src-family-kinase Lck must phosphorylate the receptor (Schoenborn et al., 2011; Tan et al., 2014). Finally, Zap70 must overcome autoinhibition before binding the phosphorylated receptor with both of its tandem SH2 domains (Deindl et al., 2009; Hsu et al., 2017). Additional proofreading steps may also contribute that are not captured in our system such as CD4/8 coreceptor scanning (Stepanek et al., 2014) and CD3ε ITAM allostery (Borroto et al., 2014). Others have also suggested receptor-level proofreading through measuring a temporal delay between receptor binding and Zap70 recruitment for both TCRs (Huse et al., 2007; Yi et al., 2019) and CARs (Bhatia et al., 2005).

We measured stronger proofreading downstream of Zap70 recruitment at the clustering of LAT and the generation of DAG. This result implies that additional kinetic proofreading steps exist separate from the receptor. The additional proofreading steps measured at LAT clustering could include the full activation of Zap70 through phosphorylation and the many phosphorylation and binding events on LAT required to initiate clustering. Hyperactive Zap70 mutants biased towards the active conformation increase T cell response to normally weak agonists, suggesting that pre-activation Zap70 short-circuits one or more proofreading steps (Shen et al., 2021).

The further proofreading measured in DAG generation suggests additional steps beyond the initial clustering of LAT. LAT does not require the phosphorylation of all its tyrosines, or binding of all known associated proteins, to cluster (Su et al., 2016). Further phosphorylation and protein recruitment events may be required for an existing LAT cluster to mature into a fully signaling-competent complex. A strong candidate for an additional proofreading step is phosphorylation of LAT Y132. Y132 is an evolutionarily-conserved poor substrate for its kinase Zap70, and this phosphorylation site is required for PLCγ recruitment, DAG generation, and Ca^++^ mobilization (Andreotti et al., 2010; Courtney et al., 2018). Mutation of the residues around Y132 to a better Zap70 substrate that is more rapidly phosphorylated resulted in increased T cell activation to normally weak agonists (Lo et al., 2019). Furthermore, Sherman et al. found that PLCγ is recruited to only a subset of LAT clusters, suggesting that only a subset of LAT clusters reach a fully signaling competent state (Sherman et al., 2011).

The kinetic proofreading model requires all intermediate steps to reset upon unbinding of the ligand (**Fig. 1A**). This means that information about the receptor’s binding state must be communicated to all proofreading steps. If kinetic proofreading steps exist beyond the T cell receptor, how is unbinding information propagated beyond the receptor? An attractive mechanism is the segregation of CD45 away from bound receptors, creating spatial regions in which activating events can occur (S. J. Davis & van der Merwe, 2006). Superresolution microscopy by Razvag et al. measured TCR/CD45 segregated regions within seconds of antigen contact at the tips of T cell microvilli (Razvag et al., 2018). Upon unbinding these regions of phosphatase exclusion collapse, and CD45 dephosphorylates receptor ITAMs and LAT clusters.

What are the advantages of distributing kinetic proofreading beyond the receptor? Having subsequent proofreading steps separate from the receptor potentially allows cooperativity between multiple receptors in activating downstream signaling steps (e.g. multiple activated receptors contributing to the creation and maturation of a single LAT cluster). Consistent with this idea, larger TCR cluster size increases the probability of T cell activation (Manz et al., 2011). Lin et al. also observed cooperativity between single TCR/pMHC binding events, where shorter binding events that overlapped in space and time activated NFAT translocation similar to long-lived single binding events in primary murine T cells (Lin et al., 2019).

Work on kinetic proofreading often assumes that the reset rate after ligand binding is uniform across all signaling intermediates (k_off_, **Fig. 1A**). However, in his seminal work applying the mathematical framework of kinetic proofreading to TCR activation, Timothy McKeithan postulated that a kinetic proofreading system with slower reset rates for downstream events compared to upstream signaling events would respond to true cognate-antigen binding events a higher percentage of the time while still ignoring weak self-antigen binding events (McKeithan, 1995). Recently, Pettman et al. modeled how slower resetting of terminal signaling complexes could account for fractional steps in their measurements of the number of proofreading steps (Pettmann et al., 2021). We measured slower dissociation of downstream LAT clusters through a slow multistep process compared to the upstream events of ligand binding and Zap-70 recruitment which followed a more rapid single step reset process (**Fig. 5C**). Our data suggests LAT clusters could survive transient unbinding events that deactivate an active receptor by being rescued by other transiently active receptors, resulting in a fully mature signaling complex created by the integration of multiple shorter binding events and potentially improving ligand discrimination as McKeithan proposed.

Cooperativity between active receptors to complete downstream proofreading steps could explain observations of T cell activation in response to intermediate affinity self-and tumour antigens (Yin et al., 2012). Intermediate lifetime binding events could last long enough to complete all necessary steps for receptor activation but not long enough to finish proofreading steps downstream of the receptor. If enough intermediate binding events overlapped in space and time, they could cooperate in accomplishing downstream proofreading events, either through the acceleration of signal propagation through the cascade or through mutual prevention of system reset upon unbinding of a subset of ligands. Such a cooperativity mechanism has been implicated in autoimmunity (Korem Kohanim et al., 2020; Wang et al., 2020) and immunotherapy off-target effects (Cameron et al., 2013).

## Methodology

Methods were adapted from our lab’s previous work (Tischer & Weiner, 2019) with modified sections highlighted below.

### Cloning

We used standard molecular biology protocols for all cloning. In general, We PCR amplified individual DNA segments and assembled them using the isothermal Gibson assembly method. Mark M Davis (Huse et al., 2007) kindly gifted us a plasmid encoding the C1 domains of PKCθ. Jay Groves kindly gifted us plasmids encoding human Zap70 and LAT (O’Donoghue et al., 2013). The Zdk-CAR was based on a CD8-CAR (Irving & Weiss, 1991), the plasmid for which was a kind gift from Art Weiss. The V529N mutation in LOV2 biases it towards the ‘open’ conformation which does not bind Zdk (Yao et al., 2008). This mutation facilitated the quick release of LOV2 from the Zdk-CAR.

### Cell culture

Jurkat cells grew in RPMI 1640 (Corning Cellgro, #10–041-CV) supplemented with 10% fetal bovine serum (Gibco, #16140–071) and glutamine (Gibco, #35050–061). We maintained Jurkats at densities between 0.1 and 1.0 × 10^6^ cells per ml. We grew 293 T cells in DMEM (Gibco, #11995–065) with 10% fetal bovine serum. All cell lines grew in humidified incubators at 37°C with 5% CO2.

### Cell line construction

We combined 1 ml of wt Jurkat cells at 0.5 × 10^6^ cells/ml with 0.5 ml lentiviral supernatant for the zdk-CAR and the appropriate reporter construct. Cells recovered overnight in the incubator, 8 ml of media was added the following day and cells were grown to desired density. Cells were labeled with the Halo dye JF549 (Grimm et al., 2015) and we sorted the desired expression levels by FACS (FACSAria II, BD). Cells recovered for approximately three passages and tested for blue-light dependent signaling on LOV2 and ICAM-1 functionalized SLBs on the microscope. We obtained wild-type Jurkat cells for this study from the laboratory of Dr. Art Weiss. Regular mycoplasma tests were negative.

### Lentiviral production

We produced lentivirus in 293 T cells using a second-generation lentiviral system (James & Vale, 2012). We transfected Cells grown to 40–60% confluency in 6-well plates with 0.5 ug each of pHR (containing the transgene of interest), pMD2.G (encoding essential packaging genes) and p8.91 (encoding VSV-G gene to pseudotype virus) using 6 ul of Trans-IT (Mirus, #MIR 2705) per manufacturer’s instructions. (Plasmids kind gift from Ron Vale.) After 48 hr, we filtered the supernatant through a 0.22 um filter and used immediately or froze at −80°C until use.

### Cell preparation for imaging

For each imaging well, we used approximately 1 × 10^6^ Jurkat cells labeled with the Halo dye JF549 (Grimm et al., 2015) (10 nM, a kind gift from the Lavis lab) for at least 15 min at 37°C before resuspension in growth media. Before imaging, we washed cells once into mHBSS-BB (at 400 RCF, 4 min), and resuspended them in 40 ul imaging media before adding them to a functionalized SLB.

### Protein purification

LOV2 and Zdk were purified and labeled as described in our previous work (Tischer & Weiner, 2019). Jay Groves generously gifted us the purified human ICAM-1-HIS used in this study (Nye & Groves, 2008).

### Glassware cleaning

All glassware was cleaned as described in our previous work (Tischer & Weiner, 2019).

### Preparation of small unilamellar vesicles (SUVs)

We washed a precleaned 4 ml glass vial 2x with chloroform (Electron Microscopy Sciences, #12550). Using Hamilton syringes (Hamilton Company, Gastight 1700 series, #80265 and #81165), we combined 1 mmoles of lipids in the following molar ratio: 97.5% DOPC, 0.5% PEG-PE, 1% DGS-NTA(Ni) and 1% biotinyl CAP PE (Avanti Polar Lipids, #850375C, #880230C, #790404C, #870282C, respectively). Next we evaporated the chloroform by slowly rotating the vial at an angle while slowly flowing nitrogen gas (Airgas, #NI 250). We vacuum desiccated the resulting lipid film for 2 hours to overnight. After desiccation, We rehydrated the lipids in 2ml of TBS and gently vortexed the vial for 10 min. We transferred the mixture to a cleaned 5 ml round bottom glass tube. We formed SUVs by submerging bottom of the tube in a Branson 1800 ultrasonic cleaner (Branson #M1800) to the level of the lipid mixture in the tube and sonicating for 30-60 minutes until clear. We added ice periodically to the sonicator bath to maintain a temperature of 0°C. Centrifugation at >21,000 RCF for 30 min at 4C (Eppendorf microcentrifuges 5425R) pelleted large lipid structures. We removed the SUV containing supernatant and used it immediately or stored it in liquid nitrogen until use.

### RCA cleaning of microscopy coverslips

We placed Glass coverslips (Ibidi, #10812) into a glass Coplin jar (Sigma-Aldrich, #BR472800) and successively bath sonicated for 10 min each in acetone (Sigma-Aldrich, #534064–4L), isopropyl alcohol (Fischer, #BP2618500), and ddH2O. Coverslips were washed five times in ddH2O between each bath sonication to remove excess organic solvents. Next we added 40 ml ddH2O, 10ml of 30% ammonium (Fischer, #423305000) hydroxide, and 10 ml 30% hydrogen peroxide (VWR, #7722-84-1) to the coverslips. We placed the Coplin jar into a 70–80°C water bath and allowed it to react for 10 min after the base solution began to vigorously bubble. We decanted the base solution and washed the coverslips five times in ddH2O. Next, we added 40ml ddH20, 10ml of 30% hydrochloric acid (Millipore Sigma, #1003180250), and 10ml 30% hydrogen peroxide to the coverslips. Again we incubated the reaction in the water bath and allowed it to react for 10 minutes after the acid solution began to vigorously bubble. We decanted the acid solution and washed the coverslips five times in ddH2O and stored them in ddH2O for up to one week.

### Functionalization of SLBs and cell preparation

After removing an RCA cleaned glass coverslip from ddH2O and immediately blown drying it with compressed nitrogen, we firmly attached a six-well Ibidi sticky chamber (Ibidi, #80608) to the coverslip. We diluted 30 ul of SUV mixture with 600 ul TBS before adding 100ul to each well and incubated at room temperature for 25 minutes. To functionalize a well, we flushed out excess lipids with 500 ul TBS. Next we added 100ul of ICAM-1-HIS diluted in TBS-BB to 150uM to the well and incubated at room temperature for 35 minutes. After incubation, we washed the well with 500 ul TBS before adding 100 ul Streptavidin (Rockland, #S000-01) diluted in TBS-BB (2 ug/ml final) to the well and incubated at room temperature for 5 min. After washing again with 500 ul TBS-BB, we added LOV2 diluted in TBS-BB (typically between 20–200 nM) to the well and incubated in the dark at 37C for 5 min. We then flushed the well with 500 ul HBSS-BB and incubated with cells previously labeled with the halo dye washed into Imaging media. Cells adhered to the SLB in the dark for 5 min at 37C before imaging on the microscope.

### Buffers for SLB functionalization and imaging

- TBS: 150mM NaCl, 20mM Tris Base, pH 7.5
- TBS-BB: TBS with 2 mg/ml BSA and 0.5 mM βME.
- mHBSS: 150mM NaCl, 40mM Kcl, 1mM CaCl2, 2mM MgCl2, 10mM Glucose, 20mM Hepes, pH 7.2
- mHBSS-BB: mHBSS with 2 mg/ml BSA and 0.5 mM βME
- Imaging media: mHBSS-BB supplemented with 2% fetal bovine serum, 50 ug/ml ascorbic acid and 1:100 dilution of ProLong Live Antifade Reagent (ThermoFisher Scientific, #P36975). Solution incubated at room temperature for at least 90 min to allow the antifade reagent to reduce oxygen levels.

### Microscopy

Imaging was performed on an Eclipse Ti inverted microscope (Nikon) with two tiers of dichroic turrets to allow simultaneous fluorescence imaging and optogenetic stimulation. The microscope was also equipped with a motorized laser TIRF illumination unit, 60x and 100x Apochromat TIRF 1.49 NA objective (Nikon), an iXon Ultra EMCCD camera (Andor), and a laser launch (Versalase, Vortran) equipped with 405-, 488-, 561-, and 640 nm laser lines. For RICM, light from a Xenon arc lamp (Lambda LS, Sutter Instrument) source was passed through a 572/35 nm excitation filter (Chroma, #ET572/35x) filter and then a 50/50 beam splitter (Chroma, #21000). Microscope and associated hardware was controlled with MicroManager (Edelstein et al., 2014) in combination with custom built Arduino controllers (Advanced Research Consulting Corporation). Blue light for optogenetic stimulation was from a 470 nm LED (Lightspeed Technologies Inc., #HPLS-36), controlled with custom micromanager scripts. For most timepoints, only RICM and TIRF561 images were collected. During and in between these timepoints, a TIRF488 long-pass dichroic mirror remained permanently in the top dichroic turret, ensuring the blue-light illumination of the cells was never interrupted. The top TIRF488 dichroic passed the longer wavelengths used for RICM and TIRF561. Only when LOV2 localization was imaged with TIRF488 at the end of a three-minute stimulation was the top dichroic removed to allow the shorter fluorescence excitation light to pass.

### Image processing

After each day of imaging, we captured TIRF488, TIRF561 and RICM images of slides with concentrated solutions of fluorescein, Rose Bengal or dPBS, respectively. To flat field correct, we subtracted the camera offset from both the experimental and dye images. By dividing the experimental image by the median dye slide image, we acquired the final flat field corrected image used in analysis (Model, 2006).

### Time course overview

We exposed cells to five minute blocks of constant blue light stimulation. Each block consisted of an initial two minute hold in strong blue light followed by a three minute stimulation at a fixed intensity of intermediate blue light. To measure the reporter output we averaged the final four timeframes of each condition. We measured CAR occupancy from a long exposure TIRF488 image taken at the end of the three minute stimulation, after the last biosensor output measurement made in TIRF561. As fluorescence excitation light from TIRF488 potently stimulates LOV2, the TIRF488 channel could only be imaged once at the very end of a three minute stimulation. We repeated five minute blocks over the course of an hour experiment with a variety of blue light intensities to sample different ligand-binding half-lives.

### CAR Occupancy Measurements

We background subtracted and background subtracted and thresholded RICM images at each timeframe to create a mask of cell footprints. The thresholded image was labeled using python skimage watershed algorithm (Van der Walt et al., 2014). A second local background mask was made by labeling pixels surrounding an expanded perimeter of each labeled cell in the cell mask. At steady state, free LOV2 should be homogeneously distributed on the SLB. The mean TIRF488 pixel intensity of a cell footprint is the sum of free LOV2 and LOV2 bound to the CAR. The mean TIRF488 pixel intensity in the background mask reflects free LOV2. Therefore, we calculated CAR occupancy as the mean TIRF488 pixel intensity in the cell mask minus the mean TIRF488 pixel intensity in the background mask.

### Biosensor measurements

To calculate biosensor output levels at steady state, we averaged the TIRF561 pixel intensity within each labeled cell mask over the last four timeframes (equivalent to the last 40s) of a three-minute hold in blue light. To account for differences in biosensor expression level, we normalized cells to their average biosensor intensity in the absence of signaling (taken as the average of the last two TIRF561 images of each two minute reset pulse of intense blue light). We sometimes observed drift in a cell’s basal activity over time. To correct for this drift, we subtracted the mean TIRF561 pixel intensity of each preceding two minute reset pulse of intense blue light from the mean value of the proceeding three-minute stimulation of blue light. Thus the reported biosensor output value is the fold-change from the cell’s average basal activity minus the fold-change from resent basal activity

Prior to model fitting, the biosensor output values of a biological replicate dataset (consisting of multiple wells acquired on the same day) were normalized by the 90th percentile output value in the dataset to properly normalize the data to the model, and to allow comparison between biosensors of variable dynamic ranges.

### Criteria for including or excluding cells in analysis

We restricted our analysis to cells that were present for the full time course. Cells that detached partway through the time course or arrived after the experiment began were excluded. Each replicate began with a no-light (maximal half-life) condition to identify non-responding cells. Cells that did not exhibit at least a 10% increase in reporter output above their basal output were excluded from analysis.

### Biological and technical replicates

Kinetic proofreading datasets (**Fig. 4**) *-* A biological replicate consisted of two time courses of stimulating cells with blue light on SLBs with different concentrations of LOV2, all on the same day (to ensure the light path did not change). We conducted each biological replicate on different days, with new preparations of cells, SLBs and LOV2. Each time course within a biological replicate contained approximately 30 cells, whose biosensor output levels and receptor occupancy were measured in all half-life conditions. As the microscopy experiments could not be done in parallel and each biological replicate took an entire day, We could not conduct technical replicates. Data underlying each replicate included in the supplement

Signaling reset datasets (**Fig. 5**) - A biological replicate consisted of four cycles of acutely terminating signaling and measuring the respective reporter lifetimes for approximately 30 cells. For each reporter, 3 or 4 biological replicates were conducted with different preparations of cells and bilayers (data included in the supplement).

### LOV2 binding half-life measurements

We calculated the average Zdk binding half-life for each blue light condition as described previously (Tischer & Weiner, 2019).

### Kinetic proofreading model fitting

We fit each biological replicate dataset to the simple model of kinetic proofreading described previously (**Fig 3D**) (Tischer & Weiner, 2019). We fit each dataset using non-linear least-squares regression as implemented by the curve_fit function from the SciPy library (Virtanen et al., 2020).

### Measuring signaling reset after ligand unbinding

OptoCAR expressing Jurkats were activated for 5 minutes on a LOV2-AF647 and ICAM-1 functionalized bilayer. At T=0, we illuminated the cell with intense blue light to unbind receptor-bound LOV2 ligands while imaging LOV2 and either Zap70 or LAT biosensors every 3 seconds. We manually segmented cells in ImageJ before tracking clusters of LOV2 and Zap70/LAT biosensors using the TrackMate plugin (Tinevez et al., 2017). We selected all tracks that persisted for at least two frames prior to blue light illumination and plotted the lifetime distribution of the resulting tracks. We then fit each lifetime distribution to erlang distributions with shape parameter k and rate parameter λ (which reduces to an exponential distribution when k=1) using curve_fit function from the SciPy library (Virtanen et al., 2020):

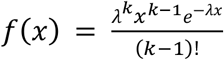

## Acknowledgments

We thank Art Weiss, Wan-Lin Lo, and Theresa Kadlecek for cell lines and thoughtful discussion, Jay Groves, Kiera Wilhelm, Mark O’Dair and Nugent Lew for purified human ICAM-1-His, expertise in SLB preparation, help with LOV2 protein purification, and thoughtful discussion. Hana El-Samad, Kirstin Meyer, Jason Town and Shohini Sen-Britain for helpful discussion and critical reading of the manuscript. This work was supported by the ARCS foundation (DB) and GM118167 (ODW), the NSF Center for Cellular Construction (DBI-1548297), and a Novo Nordisk Foundation grant for the Center for Geometrically Engineered Cellular Systems (NNF17OC0028176).

---

**Supplementary Figure 4S1.**
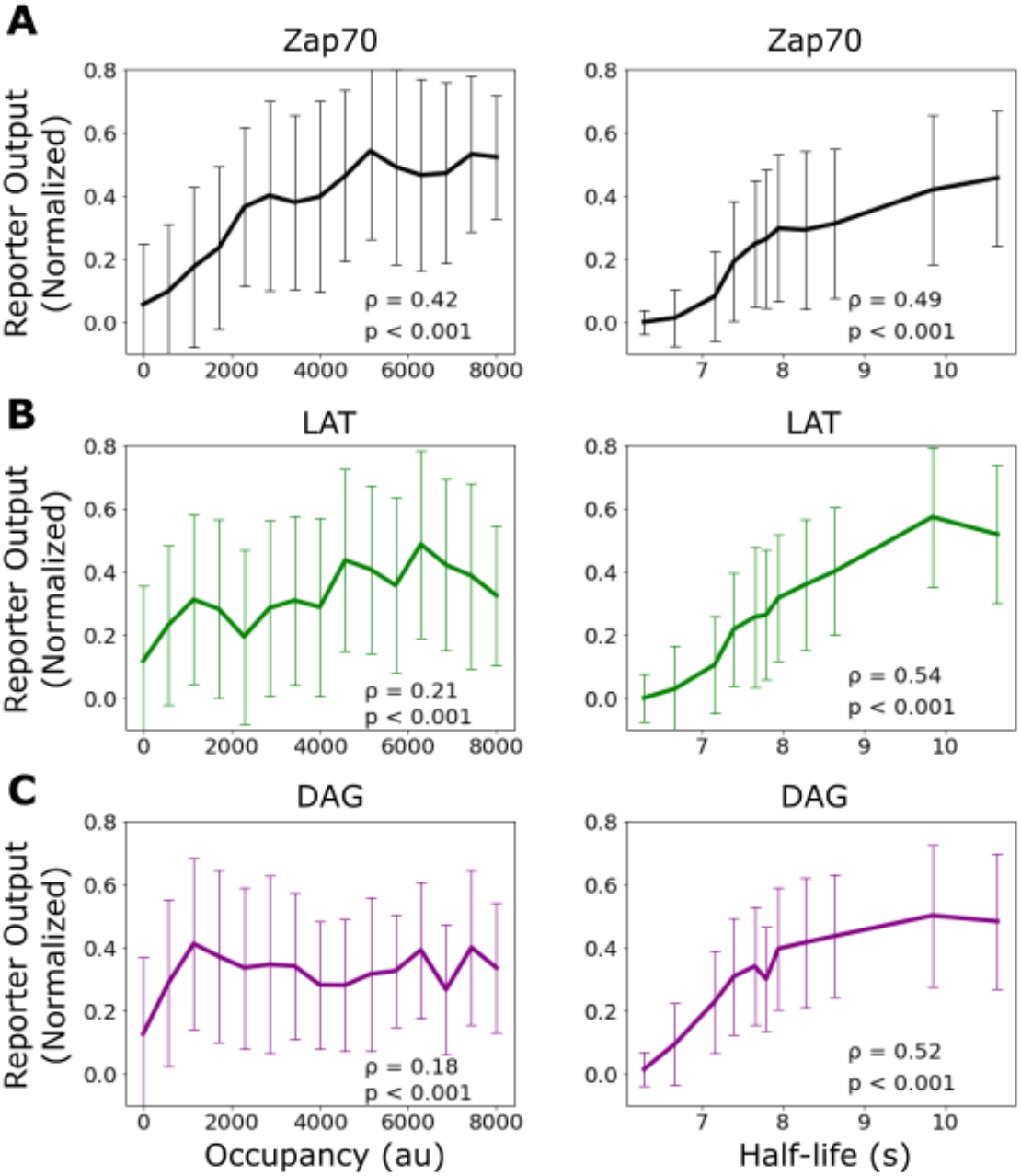
Correlation of signaling output with receptor occupancy and ligand-binding half-life. Mean reporter output (+/- 1 std) plotted as a function of receptor occupancy (left) and ligand binding half-life (right) for Zap70 recruitment **(A)**, LAT clustering **(B)**, and DAG generation **(C)**. Spearman correlation shown in the inset of each graph.

**Supplementary Figure 4S2.**
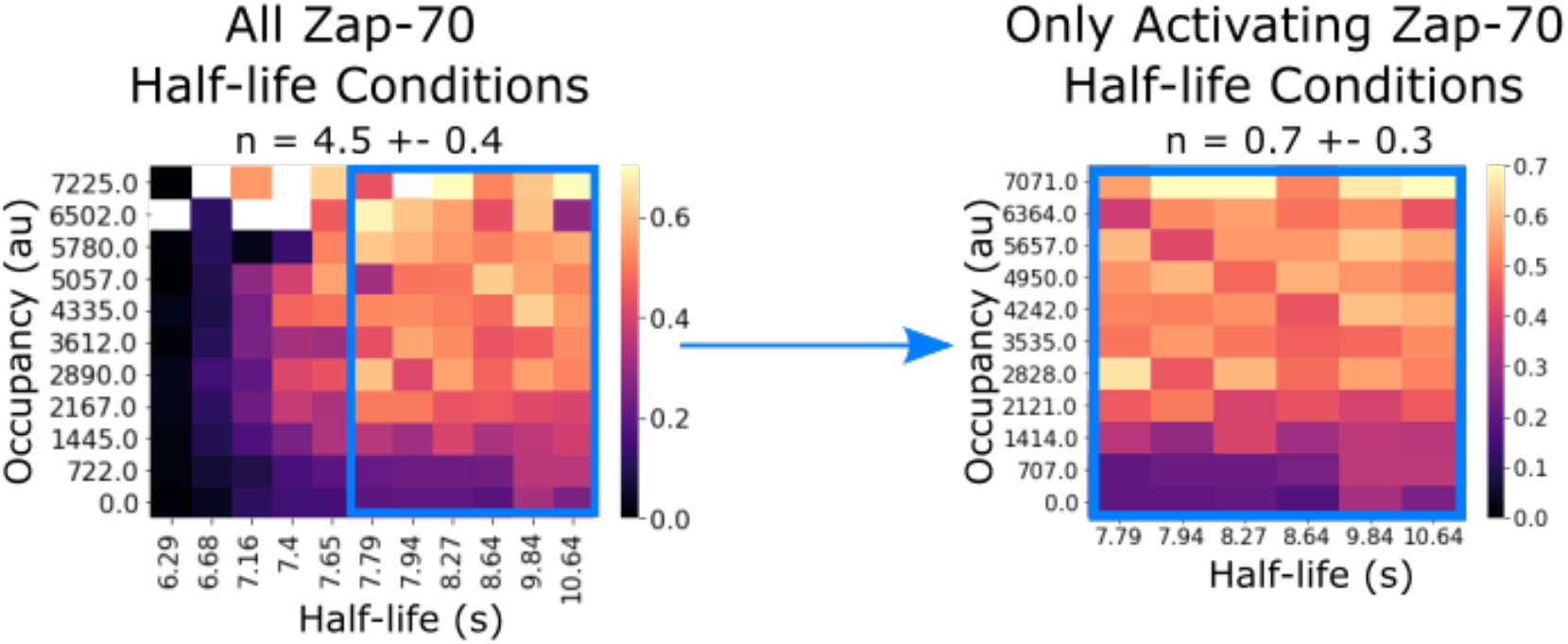
Zap70 data subset replicates previously-reported kinetic proofreading value. In our previous work, we failed to detect kinetic proofreading in Zap70 recruitment with supplementary bilayer adhesion. In our previous studies lacking light-independent bilayer adhesion, it is possible that non-activated cells did not generate sufficient footprints for segmentation. Consistent with this idea, if we restrict our analysis to only the activating half-life conditions of our current Zap70 dataset (which uses ICAM as a secondary adhesion molecule), we measure similarly low proofreading (n=0.7 +/- 0.3).

**Supplementary Figure 4S3.**
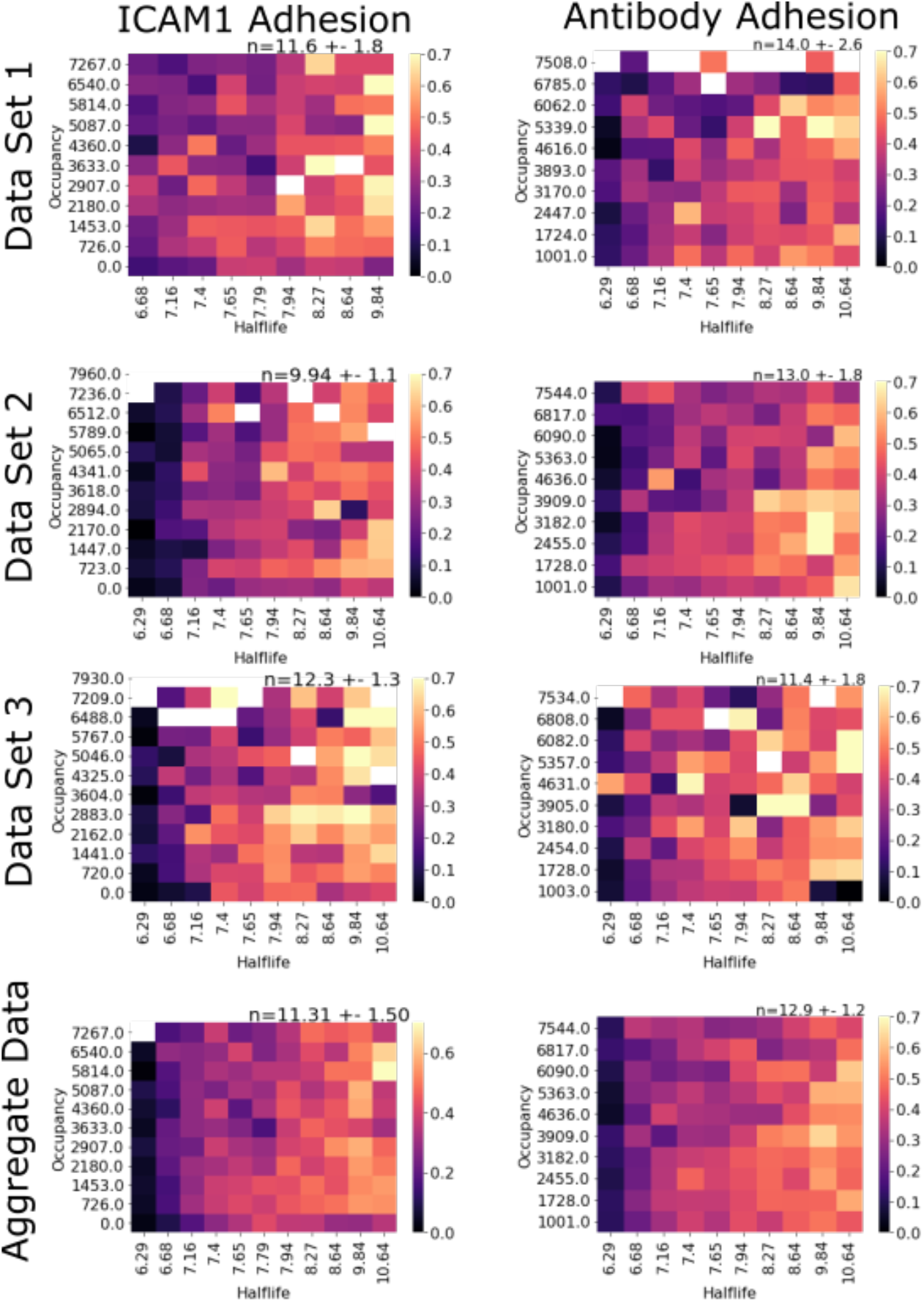
Measured kinetic proofreading at DAG generation similar for ICAM-1 and antibody-based adhesion. Bilayer adhesion through ICAM-1 or antibody produces similar kinetic proofreading model fits for the generation of DAG. Biological replicates were conducted using antibody adhesion in place of ICAM-1 and analyzed in an identical manner. Both adhesion methods fit strong proofreading to the generation of DAG.

**Supplementary Figure 5S1.**
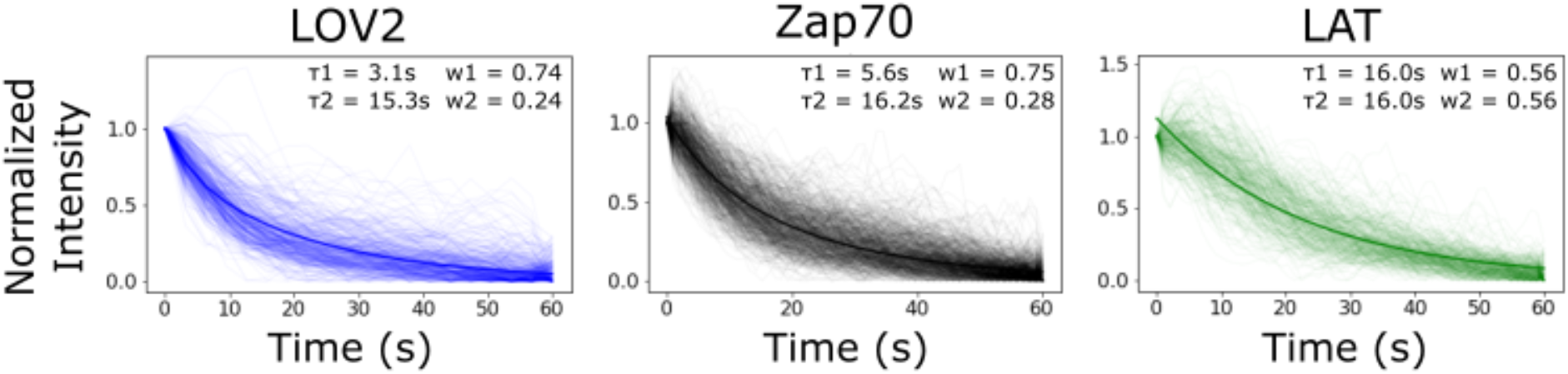
Bi-exponential fit of biosensor intensity reset curve is comparable to lifetime analysis from Figure 5. As an alternative to tracking each reporter’s lifetime distribution, we tracked the intensity of identified cluster-regions following blue light inactivation. We fit the median curve of each biosensor to a bi-exponential function. The stronger fit half-life (weight = ∼75%) for each biosensor’s bi-exponential-fit (graph inset) was comparable to the mean lifetime found analyzing the lifetime distributions (**Fig. 5C**). We found that clusters of receptor-bound LOV2 lost intensity with a half-life of 3.1 seconds. Clustered Zap70 lost intensity with a half-life of 5.6 seconds. LAT clusters lost intensity much slower than LOV2 ligand or Zap70, with a half-life of 16 seconds. The lesser fit half-life for each biosensor was approximately 16s with a weight of ∼25%. The longer half-life fit is likely a combination of photobleaching and/or cytoskeletal rearrangements of the cell occurring after receptor inactivation that are common between the biosensors.

## Supplementary Videos

**Video 1: ICAM-based adhesion yields reversible LOV2 binding for optogenetically stimulated cells**.

Time course of LOV2 on a supported lipid bilayer reversibly binding zdk-CARs expressed in Jurkat cells in the presence of ICAM-1 adhesion to the bilayer (light-independent cellular interaction). LOV2 is released from cell footprint under blue-light and rebinds in the dark (illumination status indicated in top left). Cells were adhered to the bilayer for five minutes prior to the start of the video. Two minutes of strong blue light illumination are followed by three minutes in the dark for two cycles (MM:SS time-stamp in top right).

**Video 2: ICAM-based adhesion yields reversible Zap70 recruitment for optogenetically stimulated cells**.

Time course of reversible Zap70 recruitment in Jurkat cells in the presence of ICAM-1 adhesion to the bilayer (light-independent cellular interaction). ZAP70 is recruited to phosphorylated zdk-CAR ITAMs in the dark and released back to the cell cytosol under blue-light (illumination status indicated in top left). Cells were adhered to the bilayer for five minutes prior to the start of the video. Two minutes of strong blue light illumination are followed by three minutes in the dark for two cycles (MM:SS time-stamp in top right).

**Video 3. ICAM-based adhesion yields reversible LAT clustering for optogenetically stimulated cells**.

Time course of reversible LAT clustering in Jurkat cells in the presence of ICAM-1 adhesion to the bilayer (light-independent cellular interaction). LAT forms clusters during active antigen signaling in the dark, and these clusters dissociate following blue-light-mediated dissociation of the receptor-ligand interaction (illumination status indicated in top left). Cells adhered to the bilayer for five minutes prior to the start of the video. Two minutes of strong blue light illumination are followed by three minutes in the dark for two cycles (MM:SS time-stamp in top right).

**Video 4. ICAM-based adhesion yields reversible DAG generation for optogenetically stimulated cells**.

Time course of reversible DAG generation in Jurkat cells in the presence of ICAM-1 adhesion to the bilayer (light-independent cellular interaction). DAG is generated during active antigen signaling in the dark, and DAG generation is terminated following blue-light-mediated dissociation of the receptor-ligand interaction (illumination status indicated in top left). Cells adhered to the bilayer for five minutes prior to the start of the video. Two minutes of strong blue light illumination are followed by three minutes in the dark for two cycles (MM:SS time-stamp in top right).

**Video 5: Lifetime of LOV2 and Zap70 after optogenetic signal termination**.

Example cell from time course of bound LOV2 (left) and recruited Zap70 (right) loss after inactivation of antigen signaling with strong blue light. Jurkat cells expressing zdk-CAR and Zap70 biosensor were activated on LOV2 and ICAM-1 functionalized SLBs for three minutes prior to the start of the video. Cells are illuminated with strong blue-light at the start of the video, and the lifetime of bound LOV2 and recruited Zap70 are tracked using the ImageJ TrackMate plugin (magenta circles, MM:SS timestamp top left). Only tracks originating before blue-light illumination (start of the video) were included to filter out spurious tracks later in the time course.

**Video 6. Lifetime of LOV2 and LAT after optogenetic signal termination**.

Example cell from time course of bound LOV2 (left) and LAT clusters (right) loss after inactivation of antigen signaling with strong blue light. Jurkat cells expressing zdk-CAR and LAT biosensor were activated on LOV2 and ICAM-1 functionalized SLBs for three minutes prior to the start of the video. Cells are illuminated with strong blue-light at the start of the video, and the lifetime of bound LOV2 and recruited Zap70 are tracked using the ImageJ TrackMate plugin (magenta circles, MM:SS timestamp top left). Only tracks originating before blue-light illumination (start of the video) were included to filter out spurious tracks later in the time course.

## Supplementary Files

**Figure 4 Source Data**.

Data tables (.csv) of the biological replicates used to fit the degree of kinetic proofreading (n) for each biosensor. Each row is one cell in one condition of a time course. Column “response_data” is the reporter output (zap, lat, or dag) labeled in the title of the file. Column “resonse_lov” is the occupancy measurement, and the column “half-life” is the average ligand binding half-life of the condition. We generated these data tables from raw images and plotted heatmaps using custom python scripts available at https://github.com/dbritain/KP-paper.git.

**Figure 5 Source Data**.

Data tables (.csv) for each cell used for the lifetime distributions of each reporter. Each data table is the TrackMate “spot-statistics” output of one cell over four cycles of blue-light illumination (frames 0-40, 100-140, 200-240, and 300-340, with three second interval between frames). We calculated the lifetime of TrackIDs that existed at the start of each period of blue-light illumination that did not last for the full period and fit the resulting lifetime distributions using custom python scripts available at https://github.com/dbritain/KP-paper.git.

